# Environmental and taxonomic drivers of bacterial extracellular vesicle production in marine ecosystems

**DOI:** 10.1101/2022.01.18.476865

**Authors:** Steven J. Biller, Allison Coe, Aldo A. Arellano, Keven Dooley, Jacqueline S. Gong, Emily A. Yeager, Jamie W. Becker, Sallie W. Chisholm

**Affiliations:** Department of Civil and Environmental Engineering, Massachusetts Institute of Technology, Cambridge MA; Department of Biological Sciences, Wellesley College, Wellesley MA; Department of Natural and Biomedical Sciences, Alvernia University, Reading PA; Department of Biology, Massachusetts Institute of Technology, Cambridge MA

**Keywords:** Extracellular vesicles, *Prochlorococcus*, *Pelagibacter*, *Alteromonas*, *Thalassospira*, *Alcanivorax*, *Marinobacter*, *Dokdonia*, *Polaribacter*, Station ALOHA

## Abstract

Extracellular vesicles are small (∼50-250 nm diameter) membrane-bound structures released by cells into their surrounding environment. Vesicles are abundant in the global oceans and likely play a number of ecological roles in these microbially dominated ecosystems, yet we know nothing about what influences their production and distributions. Here we examine how vesicle production varies among different strains of cultivated marine microbes and explore the degree to which this is influenced by some key environmental variables. We show that vesicle production rates – the number of vesicles produced per cell per generation – vary across an order of magnitude in cultures of marine Proteobacteria, Cyanobacteria, and Bacteroidetes. Vesicle production rates further differ among strains of the cyanobacterium *Prochlorococcus*, and vary across temperature and light gradients. These data suggest that both community composition and local environmental conditions modulate the production and standing stock of vesicles in the oceans. Examining samples from the oligotrophic North Pacific Gyre, we show depth-dependent changes in the abundance of vesicle-like particles in the upper water column in a manner broadly consistent with culture observations: highest vesicle abundances are found near the surface, where light irradiances and temperatures are greatest, and then decrease with depth. This work represents the beginnings of a quantitative framework for describing extracellular vesicle dynamics in the oceans – essential as we begin to incorporate vesicles into our ecological and biogeochemical understanding of marine ecosystems.

**Importance:** Bacteria secrete extracellular vesicles containing a wide variety of cellular compounds, including lipids, proteins, nucleic acids, and small molecules, into their surrounding environment. These structures are found in diverse microbial habitats, including the oceans, where their distributions vary throughout the water column. Differences in vesicle abundances likely affect their functional impacts within microbial ecosystems, but the factors influencing vesicle distributions in the environment remain poorly understood. Using quantitative analysis of marine microbial cultures, we show that bacterial vesicle production in the oceans is shaped by a combination of biotic and abiotic factors. Our data indicate that different marine taxa release vesicles at rates varying across an order of magnitude, and that vesicle production can change dynamically as a function of environmental conditions. Taken together with direct measurements of vesicle concentrations in the oceans, these culture-based measurements further provide a window into estimating vesicle loss rates. These findings represent a step forward in our understanding of marine vesicle distributions and provide a basis for quantitatively exploring vesicle dynamics in natural ecosystems.

## Introduction

Cells from all domains of life produce membrane-bound structures known as extracellular vesicles. Most of what we know about vesicle release by bacteria is derived from studies of model microbes relevant to advances in molecular biology or biomedical research. Recently, however, the theater has been expanded to examine the microorganisms that serve as the biogeochemical engines of natural ecosystems. In previous work, we demonstrated that the globally abundant marine cyanobacterium *Prochlorococcus* releases vesicles containing a wide range of biomolecules including lipids, nucleic acids, proteins, and metabolites (1–3). Extracellular vesicles are abundant in both coastal and oligotrophic waters, and they are produced by a diverse suite of marine bacteria, archaea, and eukaryotes from both temperate and polar regions (1, 2, 4–7). Though the precise functional roles of vesicles in marine ecosystems remain unknown, lab and field studies suggest that these structures contribute to many biological processes, including horizontal gene transfer, host-phage interactions, and nutrient exchange (1, 8–10). Vesicles represent an essentially unknown feature of the dissolved organic matter in ocean ecosystems (1). Still, we know nothing about what factors influence their production and, in turn, distributions along ocean environmental gradients.

While the production of extracellular vesicles appears to be essentially ubiquitous among bacteria, the cellular mechanisms of vesicle production remain unclear. In Gram-negative bacteria, vesicles released by intact, viable cells are largely thought to arise via ‘blebbing’ of the outer membrane, where small regions of the lipid bilayer protrude away from the cell and eventually separate in the form of a spherical vesicle (11, 12). These membrane protrusions could originate from numerous sources, such as localized membrane changes, uneven membrane synthesis, membrane turgor pressure, or the influence of small molecules on local regions of the membrane (13–17). Extracellular vesicles can arise through other mechanisms as well, such as through the pinching off of both outer and inner membrane to yield dual-membrane vesicles (18, 19), or via processes associated with the rotation of sheathed flagella (20). Self-annealing of membrane fragments released by cellular lysis can also lead to vesicle formation (21, 22).

Given the underlying complexity and diversity of vesicle biogenesis and release, understanding how this might be regulated – and even the degree to which it is directly regulated in the first place – represents a challenge. In some Gram-negative pathogens and human commensals, vesicle formation appears to involve some degree of genetic control, though it is not known whether these impacts arise through direct or indirect effects (23–28). Further, stress induced by changes in temperature (6, 29), nutrient availability, reactive oxygen species, and UV exposure has also been correlated with increased vesicle production (29–31). These examples show the potential for vesicle production to vary in response to intracellular and extracellular conditions, but the magnitude to which such variables affect vesicle release by different microbes remains an open question.

Here we examine factors that may influence vesicle production within marine microbial ecosystems, focusing on potential roles for community composition and environmental variables. To this end, we measured vesicle production rates along a spectrum of environmentally relevant conditions for a suite of taxonomically diverse, and numerically abundant, marine bacteria. We further explored the degree to which vesicle production varies among closely related strains of *Prochlorococcus*, and how vesicle production rates change across gradients of light and temperature. Finally, we examined the depth-distribution of vesicles at a site in the oligotrophic North Pacific Gyre to explore how natural vesicle abundances correlate with environmental parameters.

## Results and Discussion

### Vesicle production rates vary among marine microbial taxa

We first wondered about the degree to which marine bacteria vary in their production of extracellular vesicles. Focusing solely on vesicles released by intact, growing cells, we measured production rates by several diverse marine microbes during exponential growth in terms of vesicles produced per cell per generation, a standardized metric that enables comparisons across differences in both cell abundances and growth rates (Methods and (1)). We surveyed representatives from several of the most abundant bacterial phyla in marine systems – Proteobacteria, Cyanobacteria, and Bacteroidetes – and found that their vesicle release rates, when growing under similar conditions, spanned across an order of magnitude (Fig. 1a). Median vesicle production rates during exponential growth ranged from ∼4 vesicles cell^-1^ generation^-1^ among strains of *Prochlorococcus* to ∼58 vesicles cell^-1^ generation^-1^ in the heterotroph *Marinobacter*, with other taxa producing vesicles at intermediate values.

**FIG 1.**
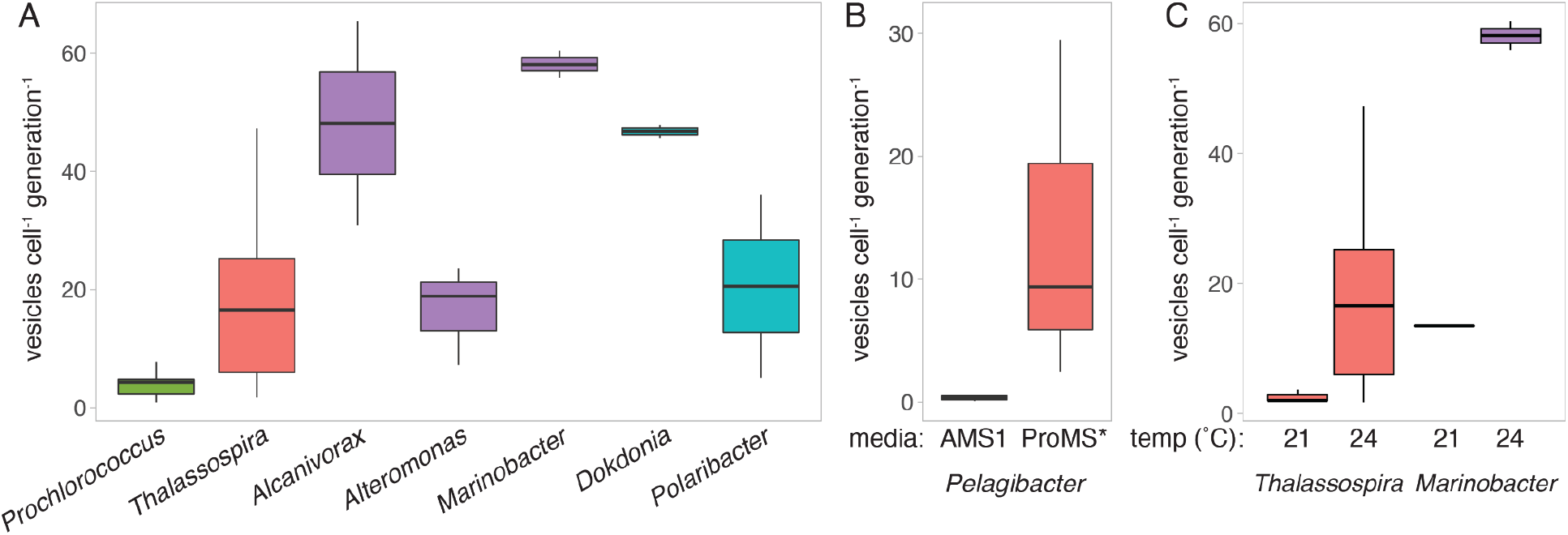
Extracellular vesicle production rates among marine microbes and as a function of growth environment. (A) Vesicle production rates for selected strains (indicated here by genus) of marine microbes at 24 °C. *Prochlorococcus* data represent median values from 5 strains grown at different light levels (see Fig. 2). (B) *Pelagibacter* (SAR11) vesicle release when grown at 22 °C on either a defined minimal medium (AMS1) or a natural seawater variant (ProMS*). (C) Impact of temperature on vesicle production by two heterotrophs. Colors represent taxonomic groupings of microbes at the Class level: Cyanophyceae (green), Alphaproteobacteria (red), Gammaproteobacteria (purple), Flavobacteriia (blue).

Though the total amount of cellular surface area available to form membrane vesicles might be expected to play some role in bounding the number of vesicles that can be formed at any time, vesicle production was not significantly associated with cell surface area (Fig. S1a). Cellular growth rates, on the other hand, were positively correlated with vesicle release rates (Pearson’s correlation *p* < 0.05), but growth rates only explained a portion of the variation in vesicle production (see below). Together, our results indicate that vesicle production rates differ markedly among marine taxa, and are due at least in part to physiological differences between strains.

### Growth conditions impact vesicle production by marine heterotrophs

To explore how environmental conditions might influence vesicle production, we first measured vesicle production rates of selected strains of marine heterotrophic bacteria under different culture conditions. Focusing first on media composition, we compared vesicle production rates by *Pelagibacter* (SAR11) grown in either a defined artificial seawater medium (AMS1) or a variant based on natural seawater but with otherwise equivalent nutrient amendments (ProMS*; see Methods). *Pelagibacter* grew 3.2x faster in ProMS* than AMS1, and vesicle release rates increased by ∼15x (from ∼0.4 vesicles cell^-1^ generation^-1^ in AMS1 to ∼6 vesicles cell^-1^ generation^-1^ in ProMS*; Fig. 1b). While we do not know which component(s) of the natural seawater background are responsible for the difference, this indicates that nutrient availability can affect vesicle production. We next asked whether changes in temperature influenced vesiculation by comparing production rates at two temperatures in *Marinobacter* and *Thalassospira*. An increase in growth temperature from 21 °C to 24 °C increased the rate of vesicle production by a factor of ∼4-8x in both strains (Fig. 1c), broadly consistent with trends observed in host-associated and terrestrial microbes such as *Pseudomonas aeruginosa* (29) and *Acinetobacter baylyi* (32).

Looking across all heterotrophs and growth conditions examined, cells tended to produce more vesicles per generation when they grew faster (Fig. S2). On average, each growth rate increase of 1 day^-1^ resulted in ∼2.6 additional vesicles released per cell per generation. However, changes in growth rate only accounted for ∼26% of the observed variance (linear regression *p* < 0.01; Fig. S2), indicating that other factors are at play. One possible explanation for the influence of growth rate on vesicle production is that increasing growth rate requires increased cell wall/membrane biosynthesis activity, which could in turn transiently weaken the connections between the outer membrane and the rest of the cell to create more opportunities for vesicle release. Environmental conditions might also modulate vesicle production via changes in membrane composition and/or fluidity (17).

### *Combinatorial impacts of light and temperature on* Prochlorococcus *vesicle production*

For abundant marine photosynthetic organisms such as *Prochlorococcus*, light and temperature are key environmental variables impacting cellular growth and physiology (33–35), and we suspected that they also influence vesicle production. To explore this, we measured production rates under different combinations of light irradiance and temperature using five different strains of *Prochlorococcus* that encompass phylogenetically and ecologically distinct high-light and low-light adapted clades (Table 1). Median production rates varied significantly among strains, and ranged from 1.2 – 7.7 vesicles cell^-1^ generation^-1^ across all conditions tested (two-way ANOVA, *p* < 0.01; Table 2; Fig. 2a). Vesicle production by individual *Prochlorococcus* strains was remarkably dynamic, varying by an average factor of ∼27 between the lowest and highest production rates observed (Table 2). On average, high light-adapted *Prochlorococcus* produced fewer vesicles per cell per generation than low light-adapted strains; the high-light strains also had a lower dynamic range of production rates (Table 2). What might account for this? High light-adapted *Prochlorococcus* are smaller than low light-adapted cells but, as with the heterotrophs, differences in cell size had little explanatory value for differences in vesicle production rates (Fig. S1b). Unlike the heterotrophs, however, differences in *Prochlorococcus* growth rate were not significantly correlated with variation in vesicle production (linear model R^2^ = 0.065; Figs. 2a and S4), suggesting that other physiological factors must play key roles.

**TABLE 1.**
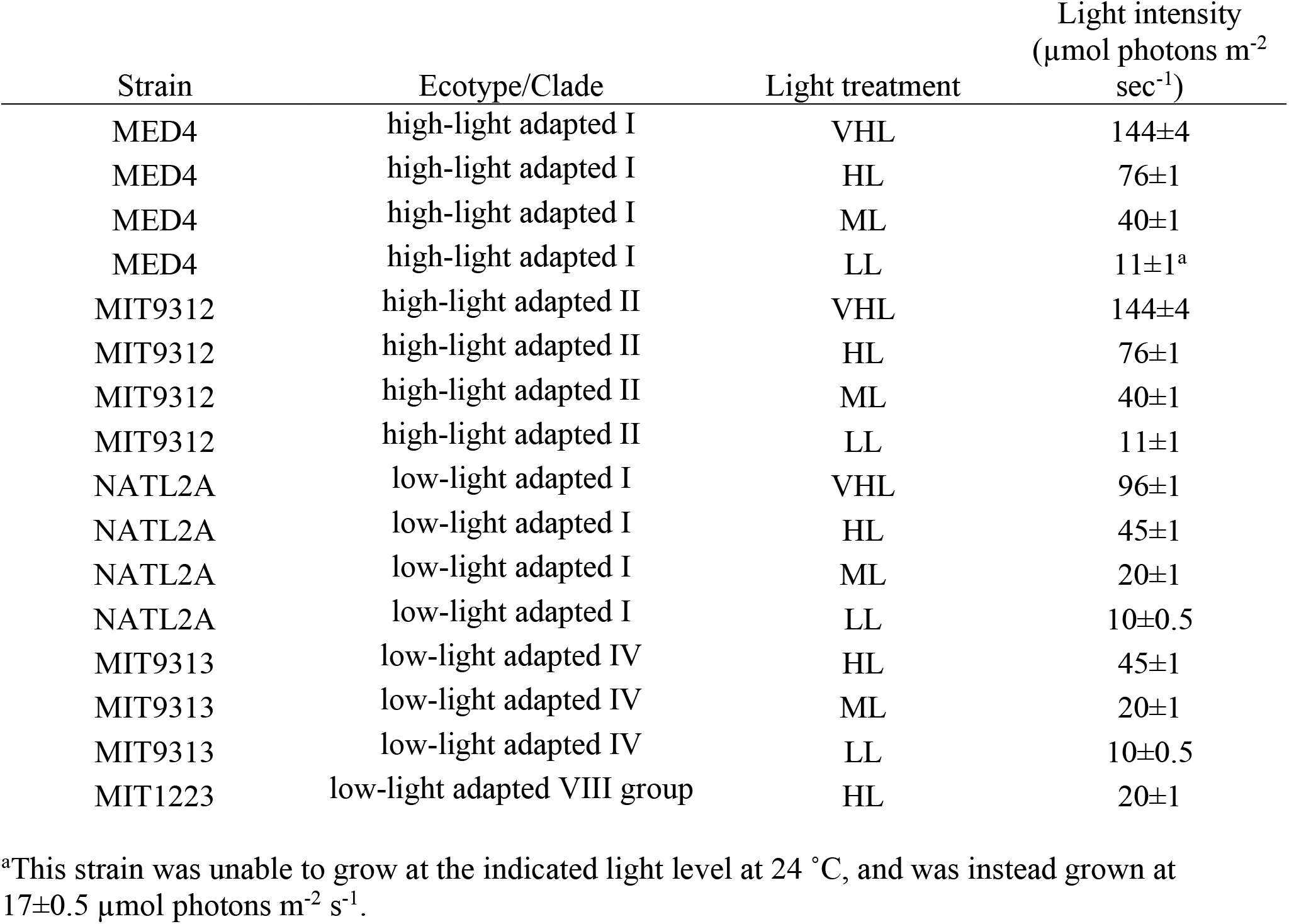
Light conditions examined for each strain. Light treatments are defined as very high light (VHL), high light (HL), medium light (ML), and low light (LL), corresponding to intensity values relative to the specific optima for that strain.

**TABLE 2.** *Prochlorococcus* vesicle production rates. All rate values are in units of vesicles cell^-1^ generation^-1^.

| Strain | Ecotype/Clade | Median vesicle production rate across all conditions tested (± SD) | Range of vesicle production rate (lowest-highest values in any combination of light/temp) | Significant variation with light (L) or temperature (T)<sup>1</sup>? |
| --- | --- | --- | --- | --- |
| MED4 | high-light adapted I | 1.2 (±1.5) | 0.2 – 3.5 | T |
| MIT9312 | high-light adapted II | 2.2 (±9.1) | 0.6 – 8.9 | - |
| NATL2A | low-light adapted I | 2.5 (±8.9) | 0.4 – 19.0 | T, L |
| MIT9313 | low-light adapted IV | 3.9 (±6.2) | 0.3 – 15.4 | T, L |
| MIT1223 | low-light adapted VIII group | 7.7 (±7.6) | 7.7 – 11.1 | *nd*<sup>2</sup> |

<sup>1</sup>Two-way ANOVA, $p < 0.05$ .
<sup>2</sup>Not determined due to limited number of conditions where this strain grew

**FIG 2.**
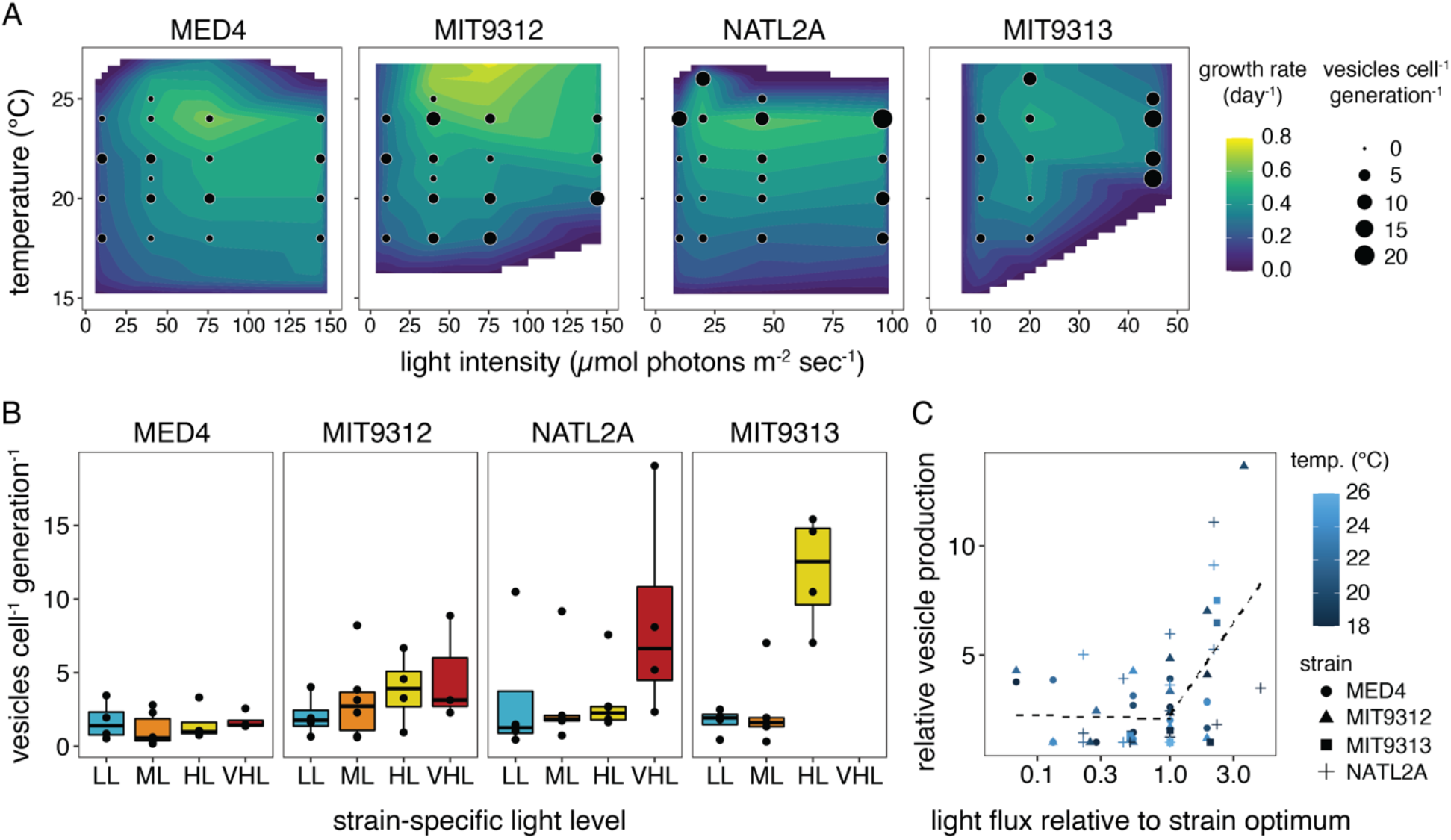
*Prochlorococcus* vesicle production rates as a function of light and temperature. (A) Vesicle production rates (black circles) for four *Prochlorococcus* strains grown at different combinations of light irradiance and temperature. Culture growth rate is indicated by background color. (B) Production rate data summarized across all growth temperatures measured. Light levels: LL= low light, ML= medium light, HL= high light, VHL=very high light (see Table 1). (C) Relative vesicle production rates as a function of relative light level. Vesicle release is normalized relative to the minimum measured value for each strain; light levels are compared to the irradiance that resulted in maximal growth rate for each strain (optimal = 1). Dashed lines represent the changepoint regression fit.

Looking at the impact of light and temperature, we found that individual strains responded differently. Vesicle production by the high-light adapted strains did not significantly change as a function of irradiance (Table 2; Fig. 2b), as consistent trends were only observed at some, but not all, temperatures (Figs. 2a, S3). By contrast, the low-light adapted cells exhibited significant variation in production rates with both light and temperature, again with higher light or increased temperature leading to increased vesicle production rates (Figs. 2a, 2b, S3; Table 2). Differences in these production patterns thus join the properties that contribute to the ecological differentiation between high-light and low-light adapted *Prochlorococcus*.

### *Vesicle release as a response to light stress in* Prochlorococcus

Our experiments indicate that light and temperature impact *Prochlorococcus* vesicle production in complex ways. Given that most strains showed an increase in production at the highest light intensities tested (Fig. 2b), we wondered whether differences in production might be associated with light stress. Irradiance levels higher than that which can be utilized by photosystems can disrupt intracellular redox balance, inhibit photosynthesis, or generate reactive oxygen species in cyanobacteria. Cells must then activate one or more stress response pathways to repair the damage and/or help to manage the excess energy input (36–38). Across all *Prochlorococcus* strains examined, production rates were not influenced by irradiance when cells were grown at or below the levels which maximized growth rates (Figs 2c, S4). Above this light optimum, when cells are assumed to experience some degree of light stress, production was significantly and positively correlated with relative increases in light (R^2^=0.23, linear model *p* < 0.01; Fig. 2c). Changepoint threshold regression analysis corroborated this difference, indicating a significant step increase in vesiculation when light levels were increased above the growth optima at each particular temperature (threshold range: 1-2x optimal light level, *p* < 0.001). This suggests that chronic light stress may directly or indirectly lead to increases in vesicle release.

Might vesicle production directly contribute to mitigating light stress, or does it instead reflect a side effect of other physiological changes occurring within the cell? Though we can only speculate at this point, cyanobacteria can utilize several strategies to dissipate excess light energy during light stress, including modifying their light-harvesting apparatus, nonphotochemical quenching, and increased production of enzymes with antioxidant functions (37). In addition, *Prochlorococcus* can utilize excess light energy to power increased levels of carbon fixation and then ‘dump’ the unneeded organic molecules into the extracellular space to maintain redox balance (36). While perhaps an indirect metabolic route, increased synthesis of lipids and secretion of fixed organic compounds via vesicles might conceivably provide yet another route for releasing the products of overflow metabolism. Intracellular oxidative stress, such as might arise from high light conditions, increases the number of vesicles produced by *Pseudomonas aeruginosa* (29) and a similar mechanism could be at play here. *Prochlorococcus* vesicles can contain oxidized metabolites, raising the possibility that they could aid the cell by removing cellular material damaged by oxidative stress (3, 12). While there remains much to uncover about this association, differences in light stress responses may explain, in part, the different degree to which light impacts vesicle release by high-light vs low-light adapted *Prochlorococcus*.

### Vesicle size varies across microbial taxa and environmental conditions

Bacteria release extracellular vesicles of varying sizes, often ranging between ∼50 - 250 nm diameter (12). Vesicle size distributions can differ among strains of the same organism (39), as a function of growth phase (40), and in response to changes in membrane composition (41). As vesicle size is associated with differences in both proteomic (42) and DNA content (43), shifts in size distributions may have functional consequences. Among the marine microbes we studied, vesicle sizes differed significantly by taxonomic group (one-way ANOVA *p* < 0.001; Fig. 3), with members of the Bacteroidetes releasing the largest (∼170 nm mode diameter) and *Prochlorococcus* releasing the smallest (∼70 nm). Vesicle size was not significantly correlated with cellular growth rate or production rate across the heterotrophs or *Prochlorococcus* strains. However, within *Prochlorococcus*, vesicle size did vary with light intensity and, to a lesser extent, growth temperature (Fig. S5; two-way ANOVA *p* < 1e-6 and *p* < 0.01, respectively). Strains grown at lower light intensities tended to release larger vesicles than cells experiencing higher light fluxes, even at levels below the optimum (Fig. S5a). The largest vesicles also tended to come from cultures grown in the middle of the temperature range tested (Fig. S5b).

**FIG 3.**
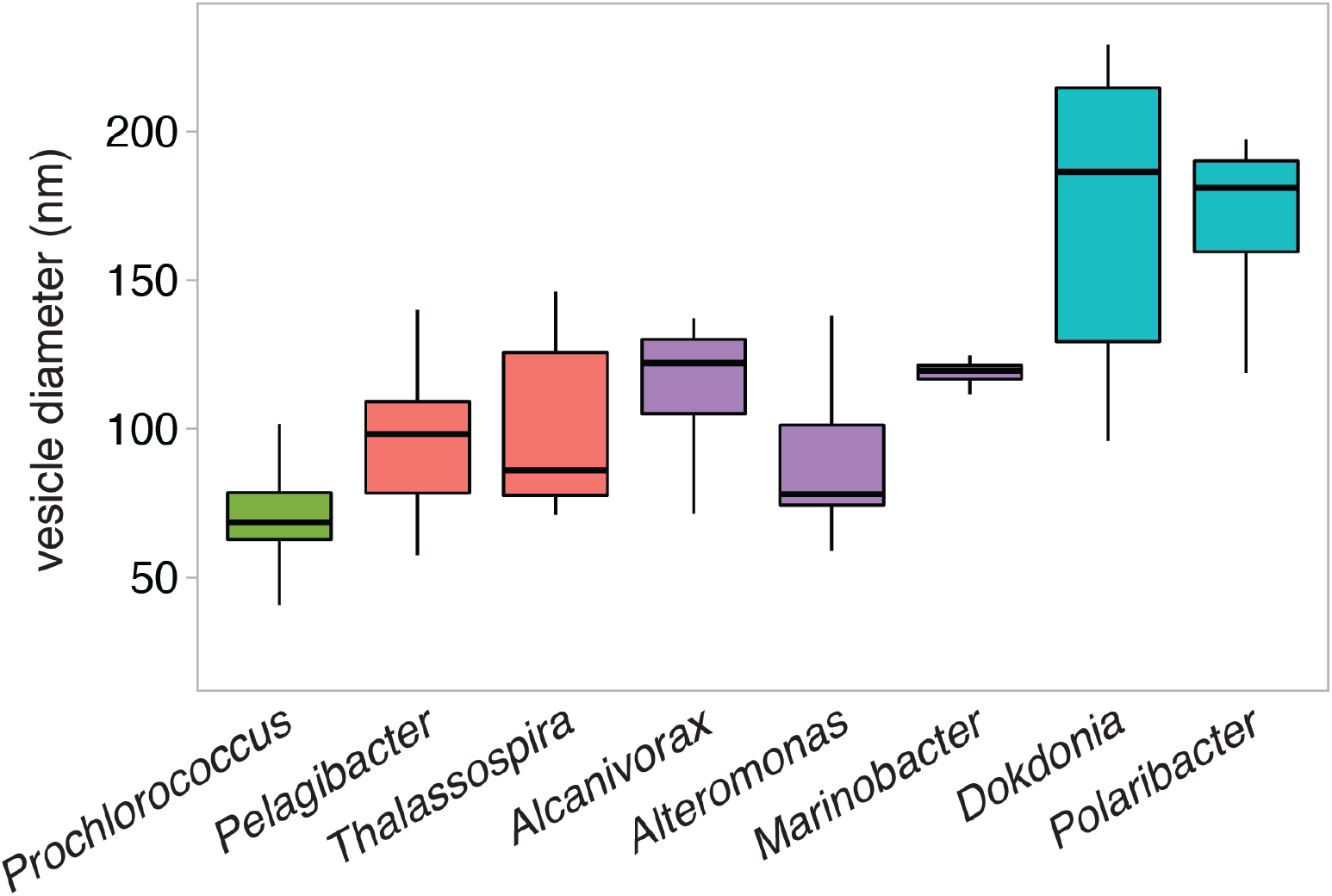
Extracellular vesicle size variation among marine microbes. Vesicle diameters indicate the mode size value from replicate vesicle populations. Colors represent taxonomic groupings of microbes at the Class level: Cyanophyceae (green), Alphaproteobacteria (red), Gammaproteobacteria (purple), Flavobacteriia (blue).

Do taxa differ in the relative investment of resources they put into vesicles? As an initial approach to addressing this question, we calculated how much membrane material is required to produce the median number of vesicles released per generation compared to each cell’s total surface area. Despite differences in cell size, vesicle diameter, and vesicle production rates, we estimate that most strains, including *Prochlorococcus, Pelagibacter*, and *Alteromonas*, release vesicles equivalent to a median of ∼3 – 6% of their cell surface lipid membranes per generation (Fig. S6). *Alcanivorax* and *Marinobacter* were notably distinct from the others and released vesicles equivalent to ∼40% of their surface area per generation (Fig. S6). Forming a given size vesicle requires proportionately less membrane material from larger cells than from smaller cells, but it appears that increased production rates can balance these lower proportional investments in individual vesicles. Given the dynamic range of vesicle production rates, and thus relative cellular investment, cells must substantially increase membrane synthesis under certain growth conditions, and this burden may not fall just on the smallest cells. The costs and benefits of changing vesicle production rates under a given environmental context, especially when considering other resources invested besides just the membranes, are clearly complex (44).

### Extracellular vesicle abundances across environmental gradients in the North Pacific

We have shown that extracellular vesicle concentrations released by the entire microbial community decreases with depth at an oligotrophic site in the Atlantic ocean (1). To explore whether those results may be generalizable to other regions of the ocean, we examined vesicle abundances at Station ALOHA in the North Pacific subtropical gyre (Fig. 4a). As at the Atlantic site, vesicle-like particles were most abundant near the surface, with their concentration decreasing by approximately an order of magnitude over the upper 500 m. Vesicle-like particle concentrations in this set of Pacific ocean samples were of the same magnitude as those previously seen in the Atlantic and declined with depth to a similar degree. This overall congruence of results across different oceans, seasons, and years, provides an early indication that this general pattern of vesicle distributions in the water column might be generally representative for mid-latitude oligotrophic waters.

**FIG 4.**
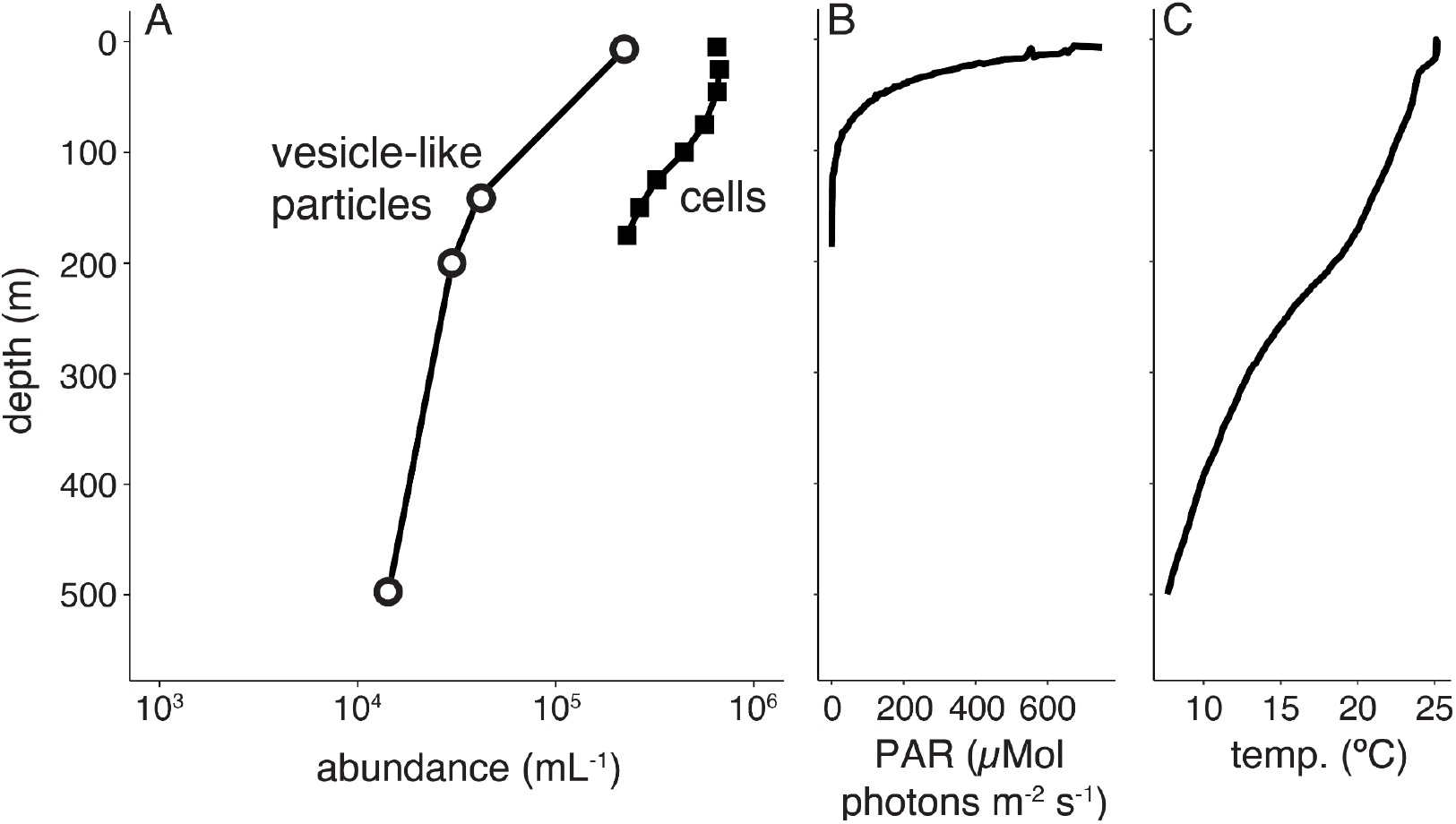
Distribution of vesicle-like particles in the North Pacific Subtropical Gyre. (A) Concentration of vesicle-like particles (open circles) and cells (filled squares) from Station ALOHA in June, 2014. Contextual environmental data are shown for (B) photosynthetically active radiation (PAR) levels and (C) temperature.

Assuming loss rates are constant with depth, the data from our culture studies would broadly predict that, all else being equal, vesicle abundances should be greatest where light irradiance and temperature are highest, and this is what we observed (Fig. 4b-c). Not only were overall vesicle-like particle concentrations greatest at the surface, but there were proportionally more vesicles per cell at the surface than in deeper samples, where environmental conditions would be expected to yield lower vesicle production rates (Fig. 4). These data are consistent with local environmental factors, such as those explored above, influencing *in situ* vesicle distribution patterns.

Given production rates in pure cultures, measurements of microbial standing stocks in the wild, and estimates of *in situ* cellular growth rates, we can also begin to estimate vesicle loss rates – an aspect of vesicle dynamics in the environment that is completely unknown. Assuming that marine phytoplankton grow at a rate of ∼0.5 per day (45, 46), with heterotrophs growing at ∼0.1 per day (45), the vesicle production rates measured here suggest that the surface community at Station ALOHA might produce in excess of 1 vesicle cell^-1^ day^-1^, or >10^5^ vesicles mL^-1^ day^-1^. This estimated daily input vesicle flux is comparable to the measured standing stocks in the oceans; if we further assume that standing vesicle concentrations are roughly constant, this would suggest that vesicle loss rates must be similar to production rates. Vesicle production rates in the ocean could certainly differ from those measured under laboratory conditions due to strain differences, nutrient availability, the influence of biotic or abiotic interactions, consideration of vesicle production through cell lysis mechanisms, or other factors. Regardless, loss rates must be significant, and could arise from processes including natural decay, consumption or degradation, fusion with other cells, or physical transport. More work is needed to arrive at a detailed estimate of losses and examine the relative contribution of these and other mechanisms.

## Conclusions

This study shows that extracellular vesicle production rates differ among marine microbes and highlights the dynamic nature of vesicle production by individual bacterial cells. The composition of extracellular vesicle populations in a given location will be influenced by a complex set of factors involving differences in cellular community structure and the vesicle production rate of each taxa under local environmental conditions. As differences in cell and vesicle concentrations would be expected to impact their encounter rates in an aqueous, planktonic environment, vesicle-mediated interaction dynamics may also changes as a result of environmental shifts. Studies addressing the fate and turnover of vesicles will be essential for developing detailed models of vesicle distributions and functions in the oceans.

## Materials and Methods

### Culturing conditions

Axenic *Prochlorococcus* cells were grown in 0.2 µm filtered and autoclaved Sargasso Sea water amended with Pro99 nutrients (47). Triplicate cultures were grown on a 14:10 hr light:dark cycle at temperatures ranging from 15 °C to 26 °C and at four light levels: very high light (VHL), high light (HL), medium light (ML), and low light (LL), with specific values determined by each strain’s light tolerance (Table 1). Culture axenicity was regularly verified using a suite of purity test broths (MPTB, ProAC, ProMM) (48–50) and by flow cytometry.

Cultures were maintained at the indicated temperature and light intensity for at least 12 generations (approximately four complete transfers), and were not sampled until the culture reached balanced growth. Growth was routinely monitored by culture fluorescence with a 10AU or TD700 Fluorometer (Turner Designs), and growth rates were calculated by exponential regression from the log-linear portion of the growth curve.

*Alcanivorax* MIT1350 was isolated from a water sample collected at 150m depth from Station ALOHA (22.75 °N 158 °W) in the North Pacific in 2013. This strain was flow-sorted from a ‘small cell, low nucleic acid’ population derived from a *Prochlorococcus* enrichment culture. Following isolation, *Alcanivorax* MIT1350, along with *Alteromonas macleodii* strain MIT1002 (51), *Thalassospira* strain MIT1004 (2), and *Marinobacter* MIT1353 (52) were maintained in ProMM medium. The genome sequence of *Alcanivorax* MIT1350 is available at the IMG database (53) under Genome ID 2681813576.

*Pelagibacterales sp*. HTCC7211, a member of the abundant warm-water surface-dwelling Ia.3 ecotype of the SAR11 bacterial clade (54), was cultured in either a defined artificial medium (AMS1; (55)) or Sargasso seawater-based ProMS* (a modified version of ProMS media (52) with pyruvate, glycine, methionine, and vitamin mixes added to concentrations equal to those in AMS1). *Dokdonia* MED134 (56) and *Polaribacter* MED152 (57) were grown in in Pro99 media supplemented with 5 g L^-1^ peptone and 1 g L^-1^ yeast extract (BD Difco). All heterotrophs were grown at 24 °C, except for HTCC7211, which was maintained at 22 °C.

We manually examined the genome sequence of each strain listed above and found no evidence for the presence of lysogenic phage and/or gene transfer agents in these organisms (based on BLASTp searches using the *Rhodobacter capsulatus* gene transfer agent sequences RCAP_rcc01682 - RCAP_rcc01698), and the cultures contained no evidence of lytic phage activity. Given this and prior electron microscopy data (2), our data assume that all particles between ∼50-250 nm diameter in these culture supernatants are extracellular vesicles.

### Cell and vesicle enumeration

At each time point sampled, cells were preserved for flow cytometry by fixation in 0.125% glutaraldehyde, flash frozen in liquid nitrogen, and stored at -80 °C until analysis. Vesicles were sampled by collecting 1 mL of culture and filtering it through a 0.22 µm Supor syringe filter (Pall). The <0.22 µm fraction was stored at -20 °C until ready for analysis. *Prochlorococcus* cells were enumerated using a Guava easyCyte 12HT flow cytometer (Luminex). Heterotrophs were stained with 1x SYBR Green I (Molecular Probes) for 1 hr and then counted using either an easyCyte 12HT or a ZE5 flow cytometer (Bio-Rad). *Prochlorococcus* cell sizes were normalized to 2 µm beads (Polysciences). Cells were excited with a blue 488 nm laser and analyzed for red fluorescence (692/40 nm), green fluorescence (530/40 nm), and size (forward scatter). Cell lengths were determined by imaging >500 cells on a Zeiss Axioskop microscope equipped with a 100x objective lens and a Nikon D90 digital camera. Cell lengths were determined using ImageJ, and calibrated using a 1mm / 0.01mm scale stage micrometer (Meiji Techno).

Vesicles were measured from samples collected at two or more timepoints during exponential growth phase. Vesicles were enumerated by nanoparticle tracking analysis using a NanoSight LM10HS instrument (Malvern/NanoSight), equipped with a LM14 blue laser module and NTA software V3.1. Three technical replicate videos (60s each) were collected from each sample using a camera level setting of 11. Samples containing >80 particles per frame were diluted with clean seawater media to a final concentration between 20-80 particles per frame. The sample chamber was thoroughly flushed with 18.2 MΩ cm^-1^ water (Milli-Q; Millipore) between samples, and visually examined to ensure that no particles were carried over between samples. Videos were analyzed using NTA V3.1 software with a threshold of 1. Vesicle concentrations in each sample are based only on particles between 50 - 250 nm diameter. Media blank controls were routinely run for each batch of media used, and any background concentration was subtracted. A small number of samples containing abnormally high particle counts (defined as containing >10x higher concentrations than other replicates of that timepoint and/or the surrounding timepoints) or in which large, highly refractive particles were present (which mask the underlying particle counts) were removed from the final dataset. The source of these contaminants is unknown and their occurrence in samples did not follow any obvious systematic patterns; we hypothesize that they could have arisen from manufacturing variation or defects in individual filters, plastic tubes used to store samples, or plastic syringes used in fluid handling.

### Vesicle production and surface area calculations

We measured vesicle production rates as the average number of vesicles produced on a per cell, per generation basis based on averaged cell and vesicle measurements from all biological replicates at each timepoint. These calculations are based on a model that incorporates the relative rates of change in cell and vesicle concentrations in pure cultures over time as detailed in (1). To better account for variation in slope across the time course, we computed the average number of vesicles produced per cell per generation between each successive pair of timepoints obtained during the exponential growth phase.

The surface area of vesicles from an individual strain was calculated assuming that vesicles are spherical structures with a diameter equivalent to the mode value measured. Cellular surface area is based on microscopy measurements and literature values (58, 59). *Prochlorococcus* was estimated to be spherical, and the surface area of other cells were estimated as cylinders with a hemisphere on each end. Relative cellular investment in vesicles per generation was defined as (individual vesicle surface area * vesicle production rate per cell per generation) / cell surface area.

### Statistics

All statistical analyses were performed in R (V 4.0.2) and plots were produced using the *ggplot2* package (60). ANOVA analysis of vesicle production rate data was performed on log-transformed values. Model selection for two-way ANOVAs was carried out using the Akaike information criterion metric, as implemented in the ‘AICcmodavg’ R package (61). Changepoint threshold regression analysis was carried out using the ‘chngpt’ R package (62). Contour plots of growth rate data were generated using the ‘metR’ and ‘interp’ R packages (63, 64).

### Field sampling

Seawater samples for vesicle analysis were collected from Station ALOHA on cruise HOT263 (June 2014). At each depth, we collected 100 - 175 L using a Niskin bottle array and concentrated the < 0.2 µm fraction by tangential flow filtration following previously described protocols (1). Vesicles were enriched across an Optiprep density gradient (Iodixanol; Sigma-Aldrich) as previously described (1). Enumeration of field concentrations were based on Nanosight concentration measurements of gradient-purified fractions containing the highest abundance of vesicles (between ∼1.14 – 1.19 g/mL) and consideration of the original sampled volume. We note that particle losses incurred during the sample processing and analysis pipeline, thus these *in situ* measurements should be considered lower-bound estimates. Cellular abundance, PAR and temperature values were provided by the Hawaii Ocean Time-series program (https://hahana.soest.hawaii.edu/hot/hot-dogs/interface.html). As vesicle-like particles could be produced by microbes from all domains of life, cell abundances represent the combined counts for cyanobacteria, heterotrophic bacteria, and eukaryotes from the indicated depths.

## Acknowledgments

We thank the captain and crew of HOT263 for their help with obtaining field samples. Stephen Giovannoni (Oregon State U.) kindly supplied *Pelagibacteriales sp*. HTCC7211, and Jarone Pinhassi (Linnaeus U.) shared the *Polaribacter* and *Dokdonia* cultures. We are also grateful to Christina Bliem and Madeline Williams for providing laboratory assistance. HOT data were obtained via the Hawaii Ocean Time-series HOT-DOGS application; University of Hawai’i at Mānoa National Science Foundation Award # 1756517.

This work was supported by the National Science Foundation (OCE-2049004 and DBI-2018337 to SJB, OCE-1356460 to SWC) and the Simons Foundation (SCOPE Award ID 329108, SWC). This is a contribution of the Simons Collaboration on Ocean Processes and Ecology (SCOPE).

The authors declare no conflicts of interest.

---

**FIG S1.**
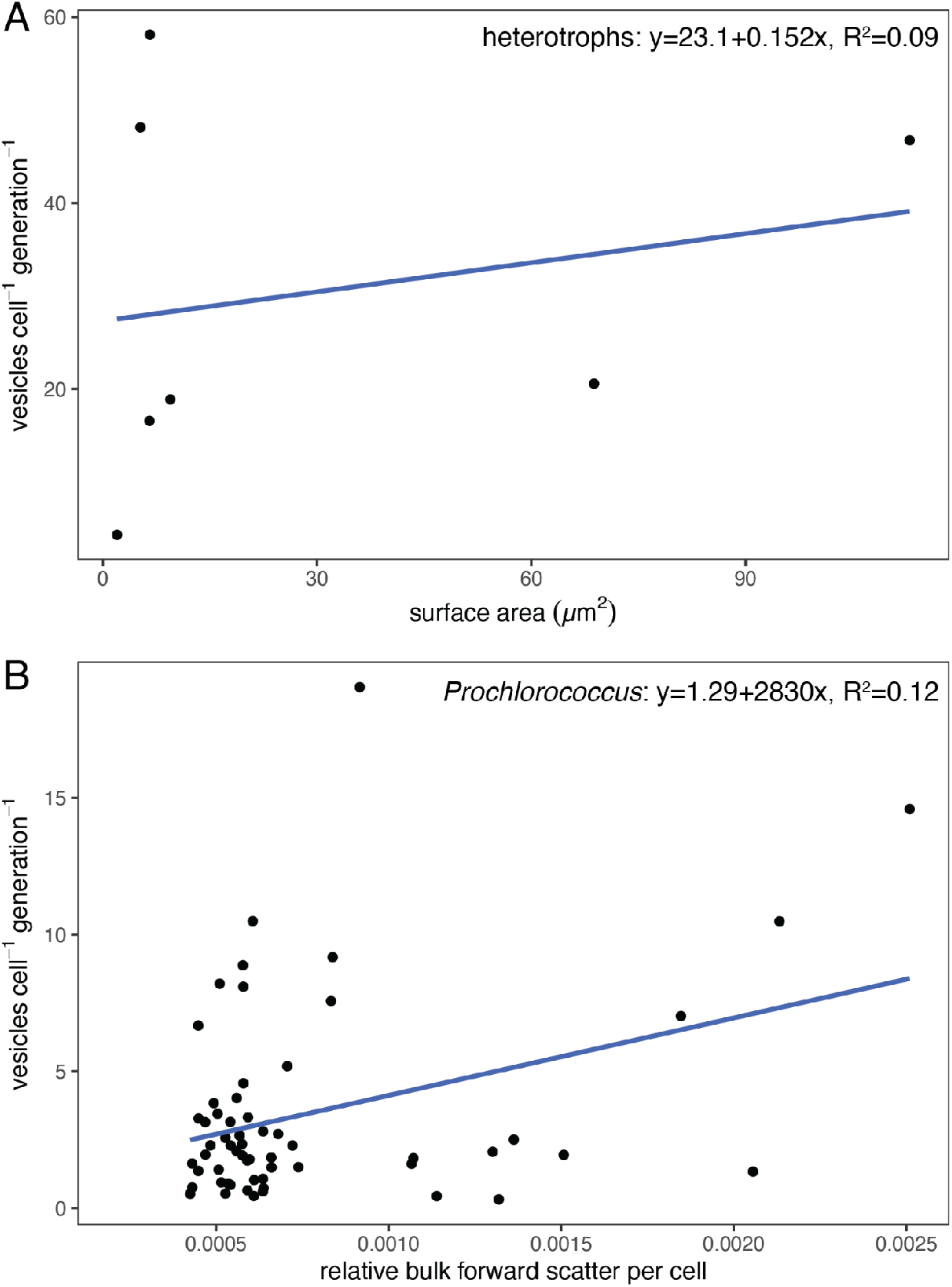
Relationship between cell size and vesicle production. (A) Cell surface area vs. median vesicle production rate across the different marine microbes shown in Fig. 1A (at 24 °C). Across all strains, no statistically significant relationship was noted (*p*=0.55). (B) Flow-cytometry based measurements of relative bulk forward light scatter (a proxy for cell size) vs. vesicle production rate specifically among *Prochlorococcus* grown at different combinations of light and temperature. A statistically significant relationship was noted (*p* < 0.01).

**FIG S2.**
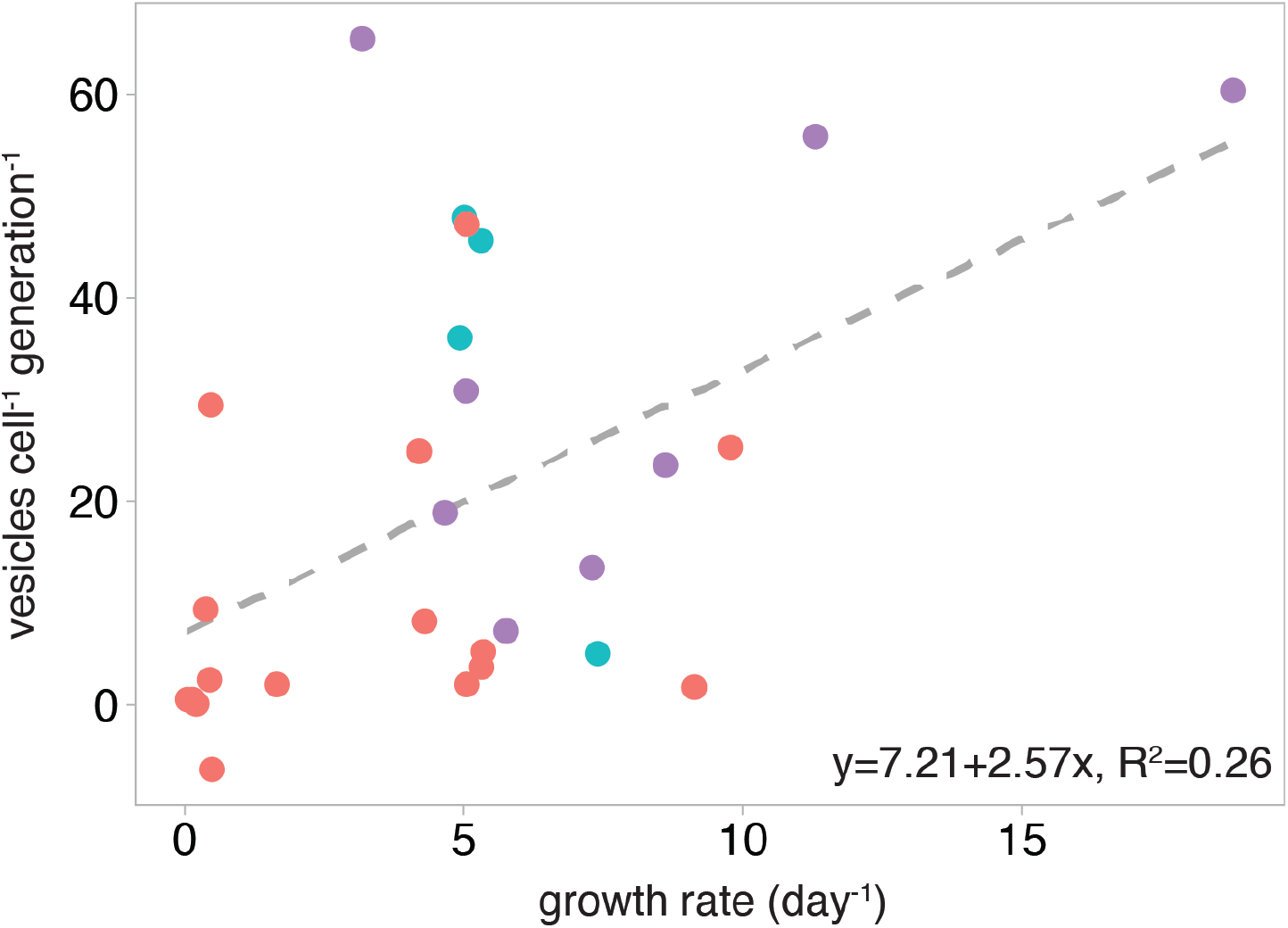
Relationship between heterotroph growth rate and vesicle production. Points indicate measured vesicle production rates across marine heterotrophs. Regression analysis was run on data from all strains and conditions shown in Fig. 1A-C. Colors represent taxonomic groupings of microbes: Alphaproteobacteria (red), Flavobacteriia (blue), Gammaproteobacteria (purple).

**FIG S3.**
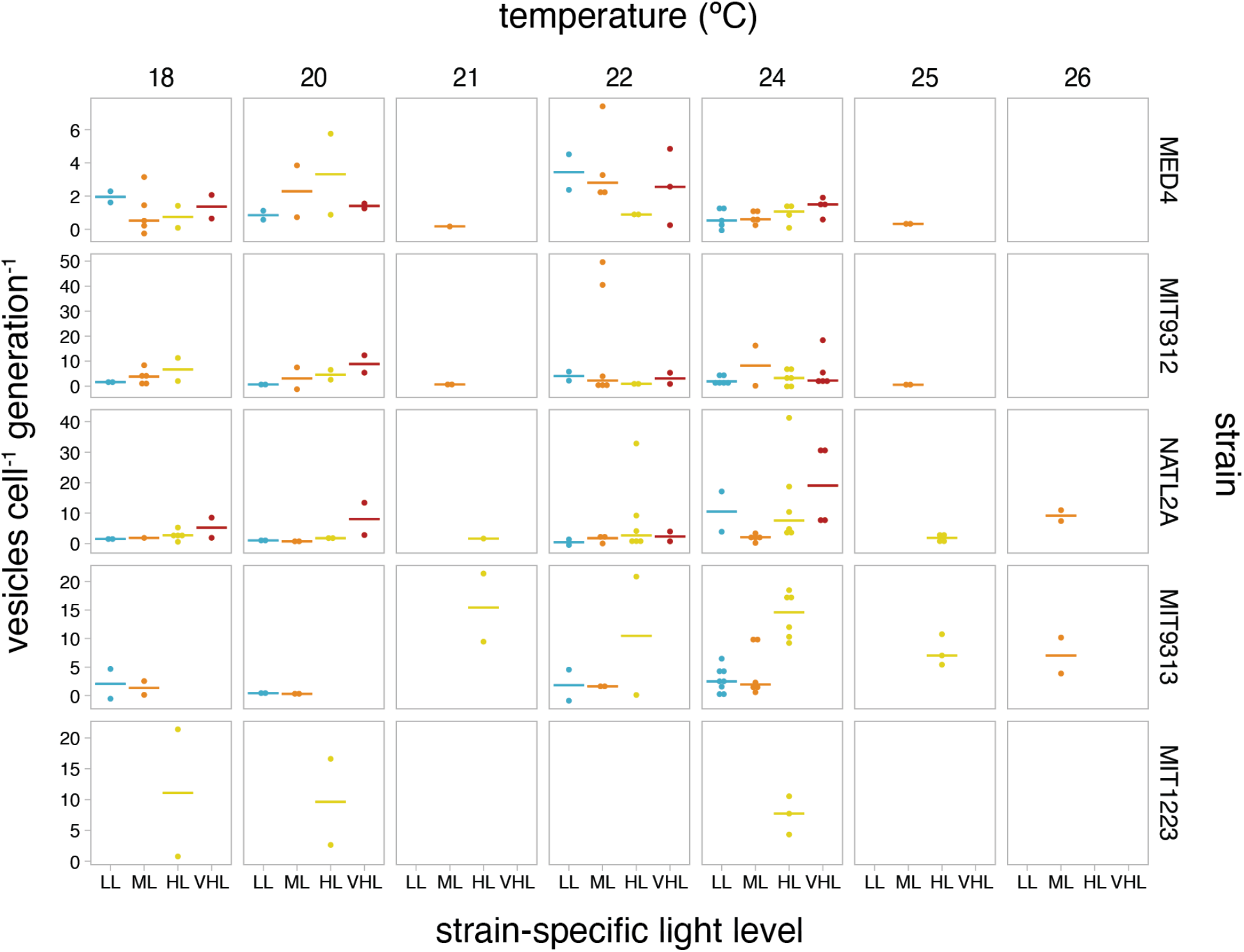
Detailed *Prochlorococcus* vesicle production across different growth conditions. Production rate data across all combinations of strains, light, and temperature. Horizontal lines indicate the median vesicle production rate measured for the indicated set of conditions. Light levels: LL= low light, ML= medium light, HL= high light, VHL=very high light (see Table 1).

**FIG S4.**
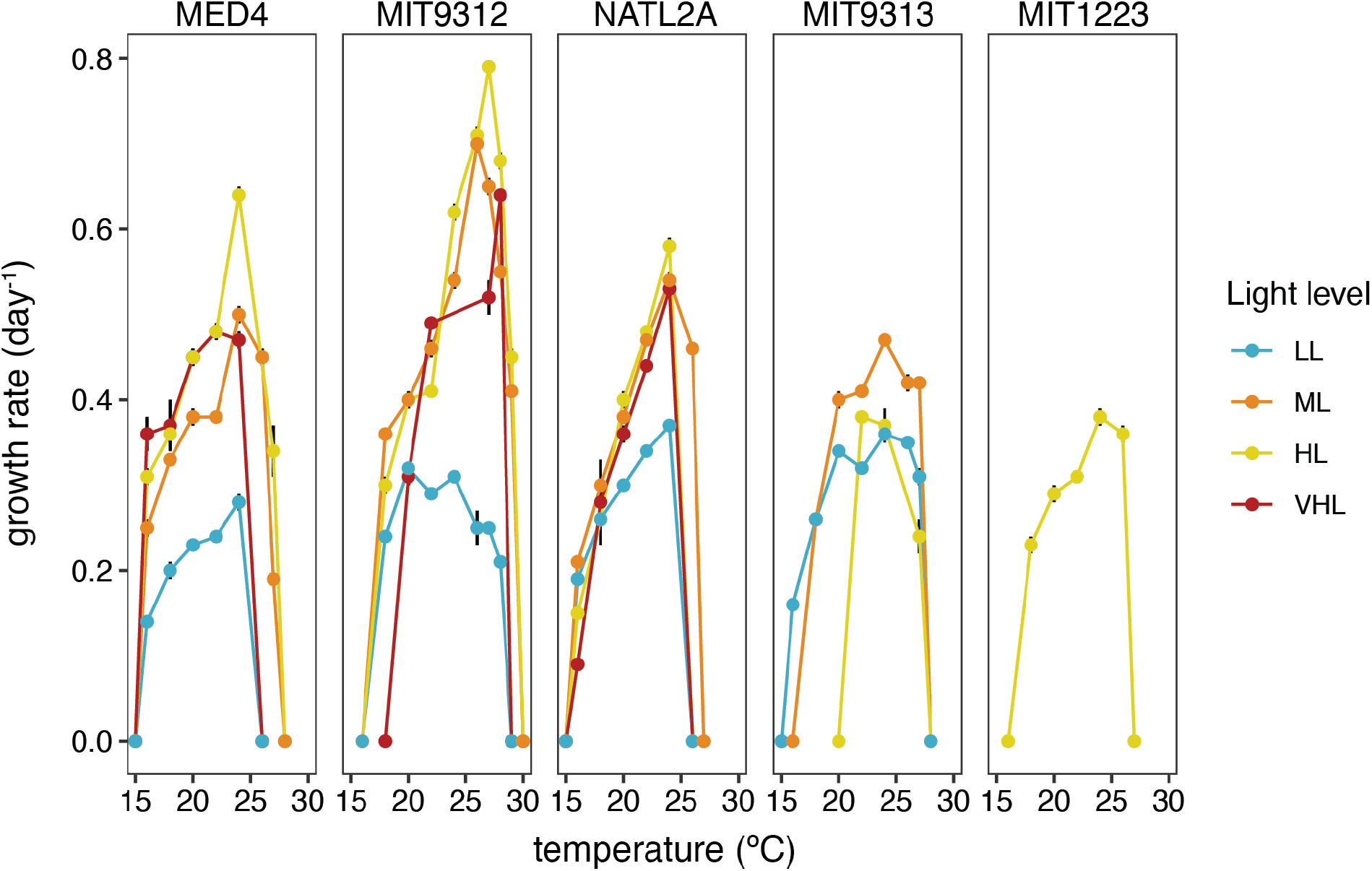
Temperature growth optima of *Prochlorococcus* strains. Values indicate measured growth rates for axenic cultures grown across different strain-specific light levels (as specified in Table 1) at the indicated temperature. Numerical values are found in Supplemental Table 1.

**FIG S5.**
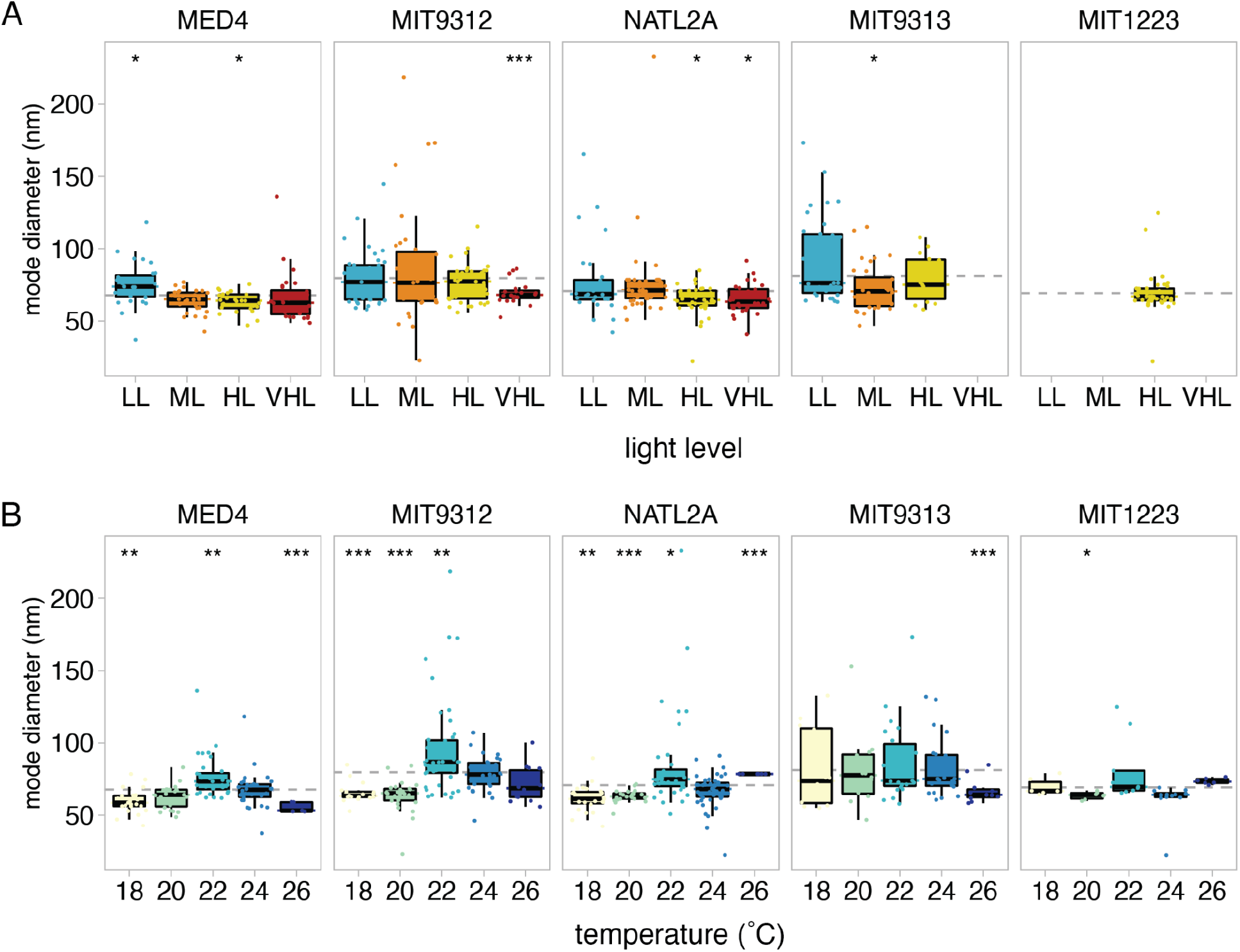
Variation in *Prochlorococcus* vesicle sizes. Values indicate the mode vesicle diameter from the indicated strains with changes in (A) light level and (B) growth temperature. Dashed grey line indicates the overall average value; asterisks indicate conditions that differ significantly from this mean (*t-*test; *: p <= 0.05, **: p <= 0.01, ***: p <= 0.001).

**FIG S6.**
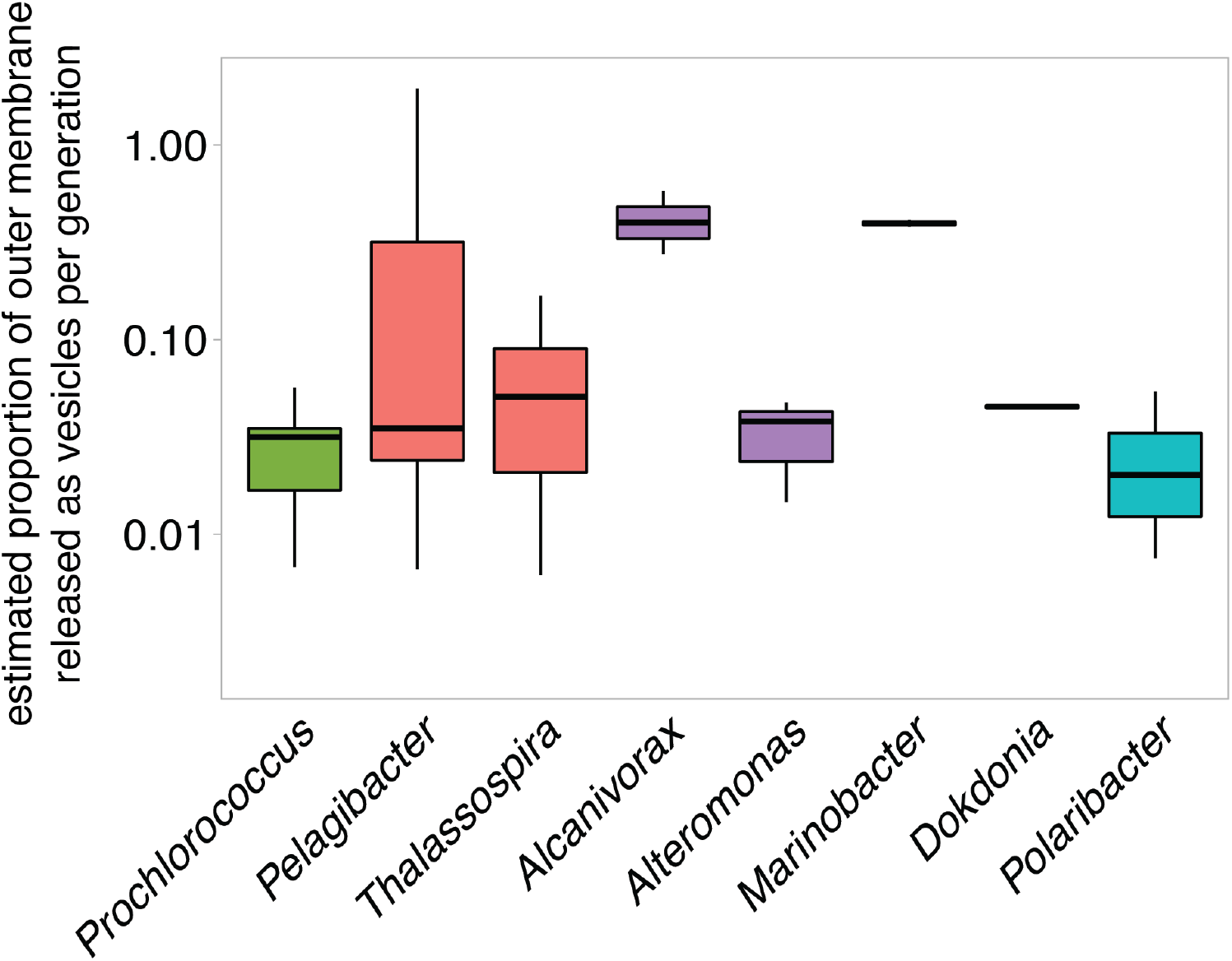
Cellular resource investment in vesicles across taxa. Relative cellular cost of vesicle production is estimated based on total vesicle surface area released per generation as a proportion of cellular surface area. Vesicle production rate ranges are based on data from Figure 1A-B. Colors represent taxonomic groupings of microbes at the Class level: Cyanophyceae (green), Alphaproteobacteria (red), Gammaproteobacteria (purple), Flavobacteriia (blue).

