## Supplemental Figures S1-S6; Table S1 for "Environmental and taxonomic drivers of bacterial extracellular vesicle production in marine ecosystems"

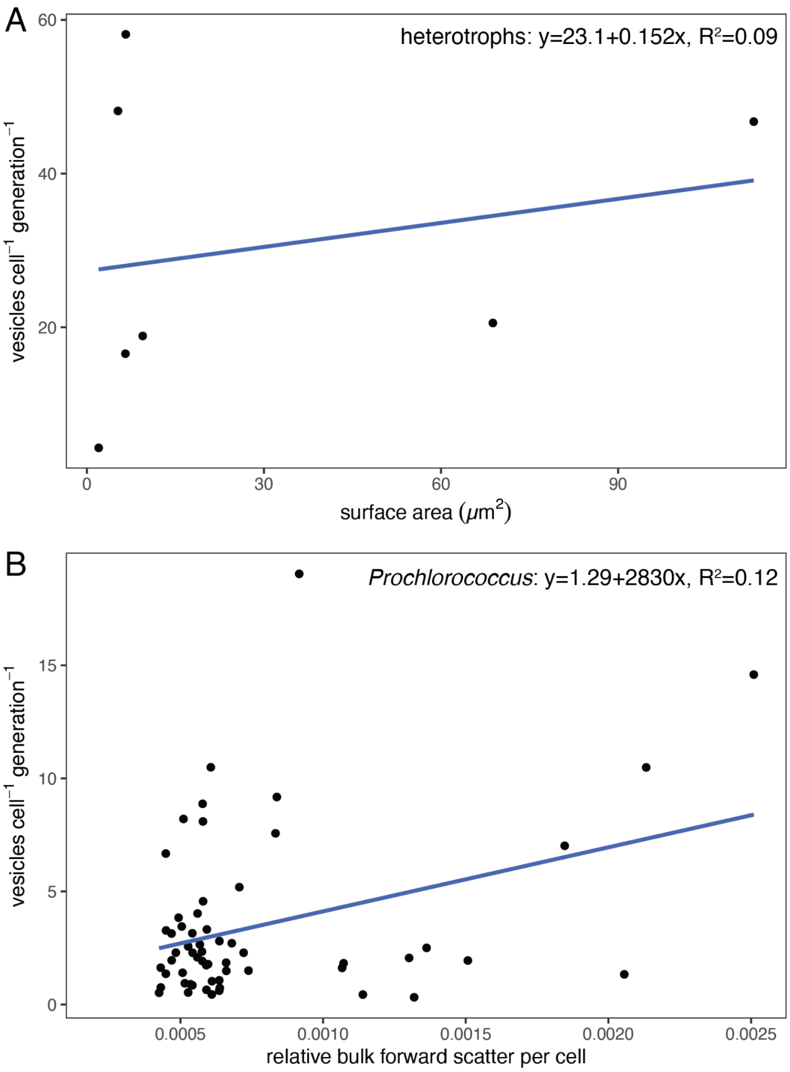


**FIG S1** Relationship between cell size and vesicle production. (A) Cell surface area vs. median vesicle production rate across the different marine microbes shown in Fig. 1A (at 24 °C). Across all strains, no statistically significant relationship was noted ($p$=0.55). (B) Flow-cytometry based measurements of relative bulk forward light scatter (a proxy for cell size) vs. vesicle production rate specifically among *Prochlorococcus* grown at different combinations of light and temperature. A statistically significant relationship was noted ($p < 0.01$).

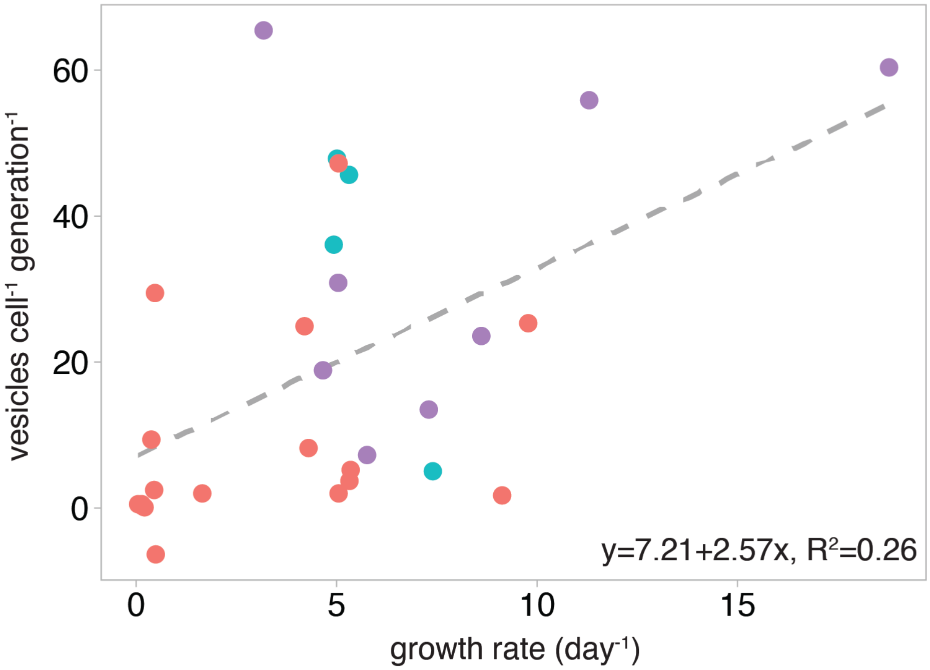


**FIG S2** Relationship between heterotroph growth rate and vesicle production. Points indicate measured vesicle production rates across marine heterotrophs. Regression analysis was run on data from all strains and conditions shown in Fig. 1A-C. Colors represent taxonomic groupings of microbes: Alphaproteobacteria (red), Flavobacteriia (blue), Gammaproteobacteria (purple).

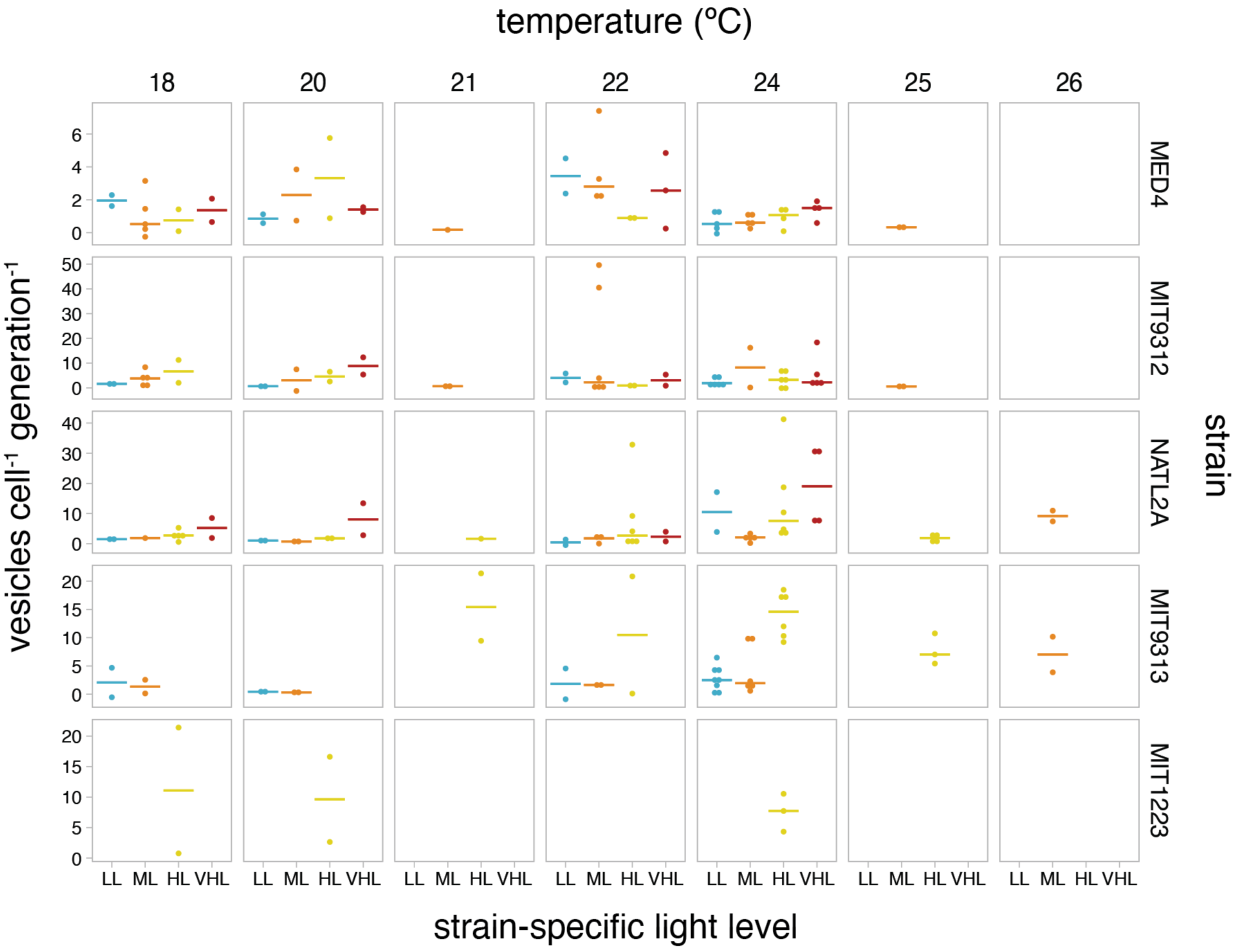


**FIG S3** Detailed *Prochlorococcus* vesicle production across different growth conditions. Production rate data across all combinations of strains, light, and temperature. Horizontal lines indicate the median vesicle production rate measured for the indicated set of conditions. Light levels: LL= low light, ML= medium light, HL= high light, VHL=very high light (see Table 1).

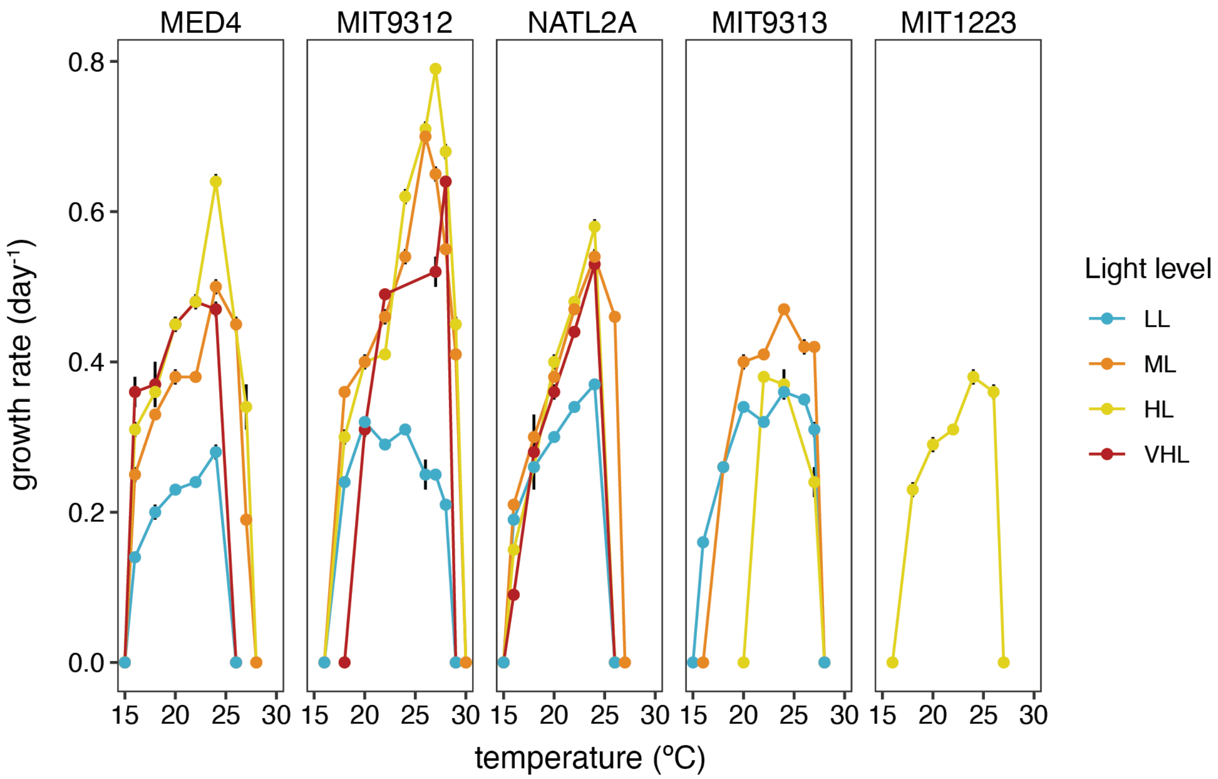


**FIG S4** Temperature growth optima of *Prochlorococcus* strains. Values indicate measured growth rates for axenic cultures grown across different strain-specific light levels (as specified in Table 1) at the indicated temperature. Numerical values are found in Supplemental Table 1.

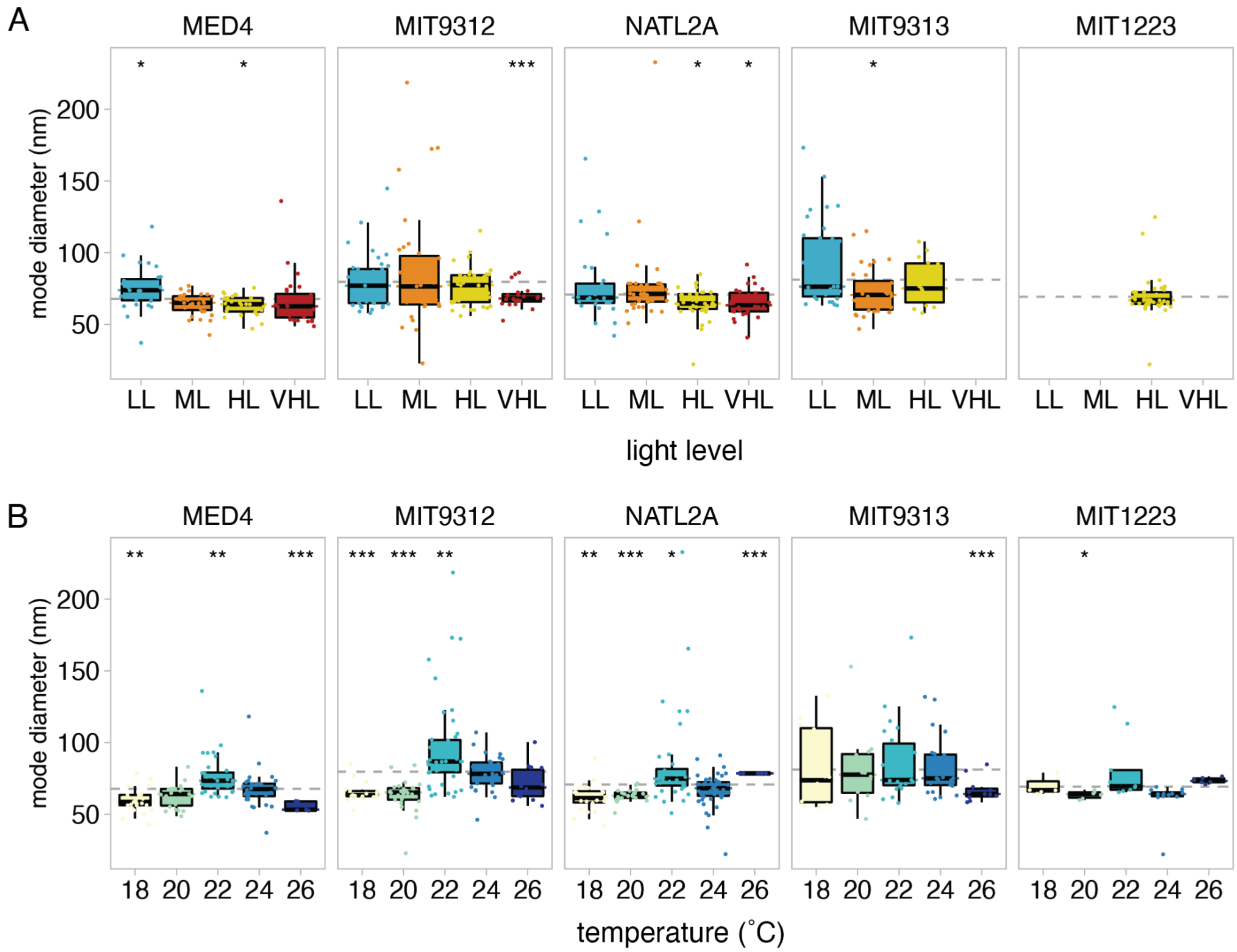


**FIG S5** Variation in *Prochlorococcus* vesicle sizes. Values indicate the mode vesicle diameter from the indicated strains with changes in (A) light level and (B) growth temperature. Dashed grey line indicates the overall average value; asterisks indicate conditions that differ significantly from this mean (*t*-test; *: p <= 0.05, **: p <= 0.01, ***: p <= 0.001).

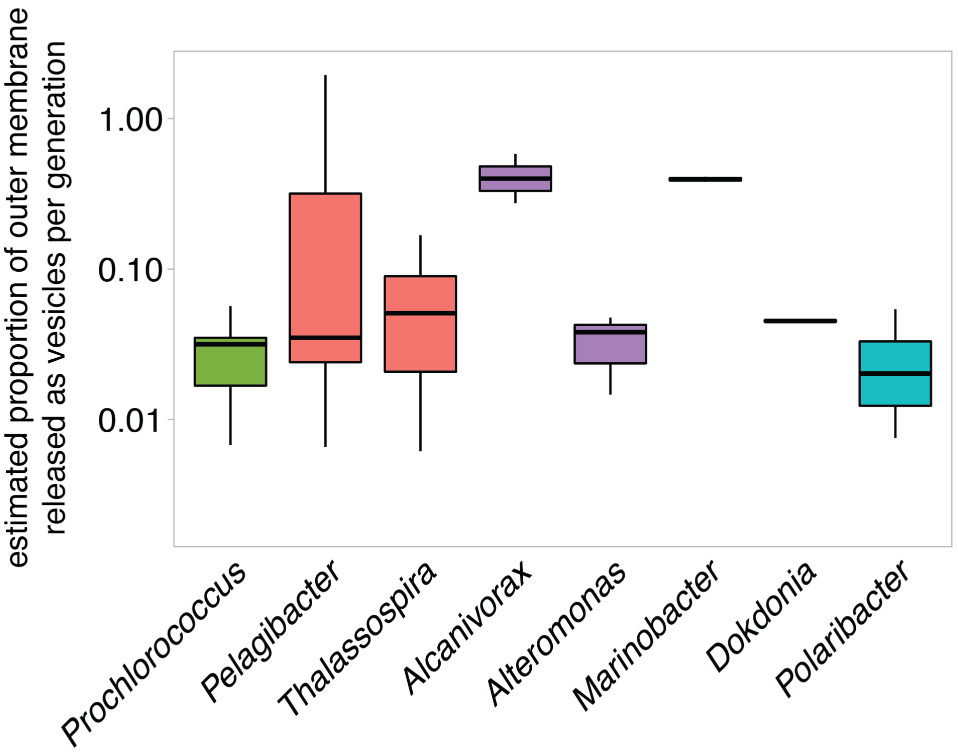


**FIG S6** Cellular resource investment in vesicles across taxa. Relative cellular cost of vesicle production is estimated based on total vesicle surface area released per generation as a proportion of cellular surface area. Vesicle production rate ranges are based on data from Figure 1A-B. Colors represent taxonomic groupings of microbes at the Class level: Cyanophyceae (green), Alphaproteobacteria (red), Gammaproteobacteria (purple), Flavobacteriia (blue).

**TABLE S1.** Exponential growth rates of axenic *Prochlorococcus* cultures (± SD) at each indicated temperature and light level.

| | | | Temperature (˚C) | | | | | | | | | | |
|---|---|---|---|---|---|---|---|---|---|---|---|---|---|
| strain | light treatment | light level (µmol photons $m^{-2}$ $sec^{-1}$) | 15 | 16 | 18 | 20 | 22 | 24 | 26 | 27 | 28 | 29 | 30 |
| MED4ax | VHL | 144 | 0.00 ± 0.00 | 0.36 ± 0.02 | 0.37 ± 0.03 | 0.45 ± 0.01 | 0.48 ± 0.00 | 0.47 ± 0.01 | 0.00 ± 0.00 | - | - | - | - |
| MED4ax | HL | 76 | 0.00 ± 0.00 | 0.31 ± 0.01 | 0.36 ± 0.01 | 0.45 ± 0.01 | 0.48 ± 0.01 | 0.64 ± 0.01 | - | 0.34 ± 0.03 | 0.00 ± 0.00 | - | - |
| MED4ax | ML | 40 | 0.00 ± 0.00 | 0.25 ± 0.01 | 0.33 ± 0.00 | 0.38 ± 0.01 | 0.38 ± 0.00 | 0.50 ± 0.01 | 0.45 ± 0.01 | 0.19 ± 0.00 | 0.00 ± 0.00 | - | - |
| MED4ax | LL | 10 | 0.00 ± 0.00 | 0.14 ± 0.00 | 0.20 ± 0.01 | 0.23 ± 0.00 | 0.24 ± 0.00 | 0.28 ± 0.01 | 0.00 ± 0.00 | - | - | - | - |
| MIT9312ax | VHL | 144 | - | - | 0.00 ± 0.00 | 0.31 ± 0.01 | 0.49 ± 0.00 | - | - | 0.52 ± 0.02 | 0.64 ± 0.00 | 0.00 ± 0.00 | - |
| MIT9312ax | HL | 76 | - | 0.00 ± 0.00 | 0.30 ± 0.01 | 0.40 ± 0.01 | 0.41 ± 0.00 | 0.62 ± 0.01 | 0.71 ± 0.01 | 0.79 ± 0.00 | 0.68 ± 0.01 | 0.45 ± 0.01 | 0.00 ± 0.00 |
| MIT9312ax | ML | 40 | - | 0.00 ± 0.00 | 0.36 ± 0.00 | 0.40 ± 0.00 | 0.46 ± 0.01 | 0.54 ± 0.01 | 0.70 ± 0.01 | 0.65 ± 0.01 | 0.55 ± 0.00 | 0.41 ± 0.00 | 0.00 ± 0.00 |
| MIT9312ax | LL | 10 | - | 0.00 ± 0.00 | 0.24 ± 0.00 | 0.32 ± 0.00 | 0.29 ± 0.00 | 0.31 ± 0.00 | 0.25 ± 0.02 | 0.25 ± 0.00 | 0.21 ± 0.00 | 0.00 ± 0.00 | - |
| NATL2Aax | VHL | 96 | 0.00 ± 0.00 | 0.09 ± 0.00 | 0.28 ± 0.05 | 0.36 ± 0.01 | 0.44 ± 0.00 | 0.53 ± 0.01 | 0.00 ± 0.00 | - | - | - | - |
| NATL2Aax | HL | 45 | 0.00 ± 0.00 | 0.15 ± 0.00 | 0.26 ± 0.00 | 0.40 ± 0.01 | 0.48 ± 0.00 | 0.58 ± 0.01 | 0.00 ± 0.00 | - | - | - | - |
| NATL2Aax | ML | 20 | 0.00 ± 0.00 | 0.21 ± 0.00 | 0.30 ± 0.00 | 0.38 ± 0.00 | 0.47 ± 0.00 | 0.54 ± 0.01 | 0.46 ± 0.00 | 0.00 ± 0.00 | - | - | - |
| NATL2Aax | LL | 10 | 0.00 ± 0.00 | 0.19 ± 0.00 | 0.26 ± 0.01 | 0.30 ± 0.00 | 0.34 ± 0.00 | 0.37 ± 0.00 | 0.00 ± 0.00 | - | - | - | - |
| MIT9313ax | HL | 45 | - | - | - | 0.00 ± 0.00 | 0.38 ± 0.00 | 0.37 ± 0.02 | - | 0.24 ± 0.02 | 0.00 ± 0.00 | - | - |
| MIT9313ax | ML | 20 | - | 0.00 ± 0.00 | 0.26 ± 0.00 | 0.40 ± 0.01 | 0.41 ± 0.00 | 0.47 ± 0.00 | 0.42 ± 0.01 | 0.42 ± 0.00 | 0.00 ± 0.00 | - | - |
| MIT9313ax | LL | 10 | 0.00 ± 0.00 | 0.16 ± 0.00 | 0.26 ± 0.00 | 0.34 ± 0.00 | 0.32 ± 0.00 | 0.36 ± 0.00 | 0.35 ± 0.00 | 0.31 ± 0.01 | 0.00 ± 0.00 | - | - |
